# Relative timing and coupling of neural population bursts in large-scale recordings from multiple neuron populations

**DOI:** 10.1101/2025.02.18.638950

**Authors:** Motolani Olarinre, Joshua H. Siegle, Robert E. Kass

## Abstract

The onset of a sensory stimulus elicits transient bursts of activity in neural populations, which are presumed to convey information about the stimulus to downstream populations. The time at which these synchronized bursts reach their peak is highly variable across stimulus presentations, but the relative timing of bursts across interconnected brain regions may be less variable, especially for regions that are strongly functionally coupled. We developed a simple analytical framework that obtains good estimates of population burst times on a trial-by-trial basis, and of the correlations in the timing of evoked population bursts across areas. We show that this method performs well on simulated data, and is 85 to 90% faster than an alternative, recently-published method while also being much easier to apply. Using this new approach, we examined the relative timing of the first two population bursts following the onset of a drifting grating stimulus in large-scale recordings of spiking activity from six cortical visual areas and one visual thalamic nucleus in thirteen mice. The new method allowed us to identify mouse-to-mouse variation in peak times and region-to-region functional coupling. While all results were consistent with known anatomy and physiology, we found some sequences of activity across areas to be the same across all mice, while others varied with the individual. The general approach can thus produce sensitive analyses of timing relationships across neural populations.

**Significant Statement:** Careful analysis can reveal strong and precisely-timed interactions across multiple brain areas from small populations of spiking neurons. We developed a computationally efficient procedure that allowed us to examine the relative timing and coupling of 7 visual areas (6 cortical and one thalamic) and compare results in over 10 mice. The method can be used to track the flow of information across the brain in response to stimuli or during a behavioral task.

## Introduction

Within tens of milliseconds after the onset of a sensory stimulus, spikes are conveyed from the periphery and evoke a transient burst of activity across large populations of neurons in the thalamus and cortex. Responses to stimulus onset in sensory cortex often include two activity peaks, with the earlier peak reflecting the feed-forward propagation of spikes from the periphery, while the second peak (occurring 100-200 ms later) likely reflects feedback from other cortical areas Sachidhanandam et al. (2013); Manita et al. (2015). The standard method for measuring the timing of these peaks is to compute a peri-stimulus time histogram (PSTH), which averages the evoked response across trials, but this ignores trial-to-trial variability in peak times, effectively discarding useful information that might give insights into the propagation of spikes through cortical areas Chen et al. (2022).

Modern electrophysiological recording techniques, such as Neuropixels probes Jun et al. (2017); Steinmetz et al. (2021), have enabled simultaneous recordings of spike trains from hundreds of neurons in multiple cortical regions, making it possible to observe the timing of evoked responses in greater detail than was previously possible Siegle et al. (2021); Jia et al. (2022). The Allen Brain Observatory Neuropixels Visual Coding Dataset Allen Institute MindScope Program (2019), an open dataset consisting of electrophysiological recordings from multiple cortical and thalamic visual areas in parallel, is a prime example of what can achieved with these probes (Figure 1A-C). The cortical areas recorded in this dataset display a stereotypical dual-peaked response to the onset of a full-field drifting grating stimulus, with the average timing of the first peaks consistent with their relative hierarchical ordering determined by anatomy (Figure 1D) Harris et al. (2019); Siegle et al. (2021); D’Souza et al. (2022). Because each peak results from the synchronous firing of many neurons in a given region, we refer to these peaks as “population bursts.” As noted by Kass et al. (2023), assuming that behaviorally relevant information is transmitted across parts of the brain through such transient bursts of activity in neural populations, their timing on a trial-by-trial basis should reveal coordinated activity.

**Figure 1:**
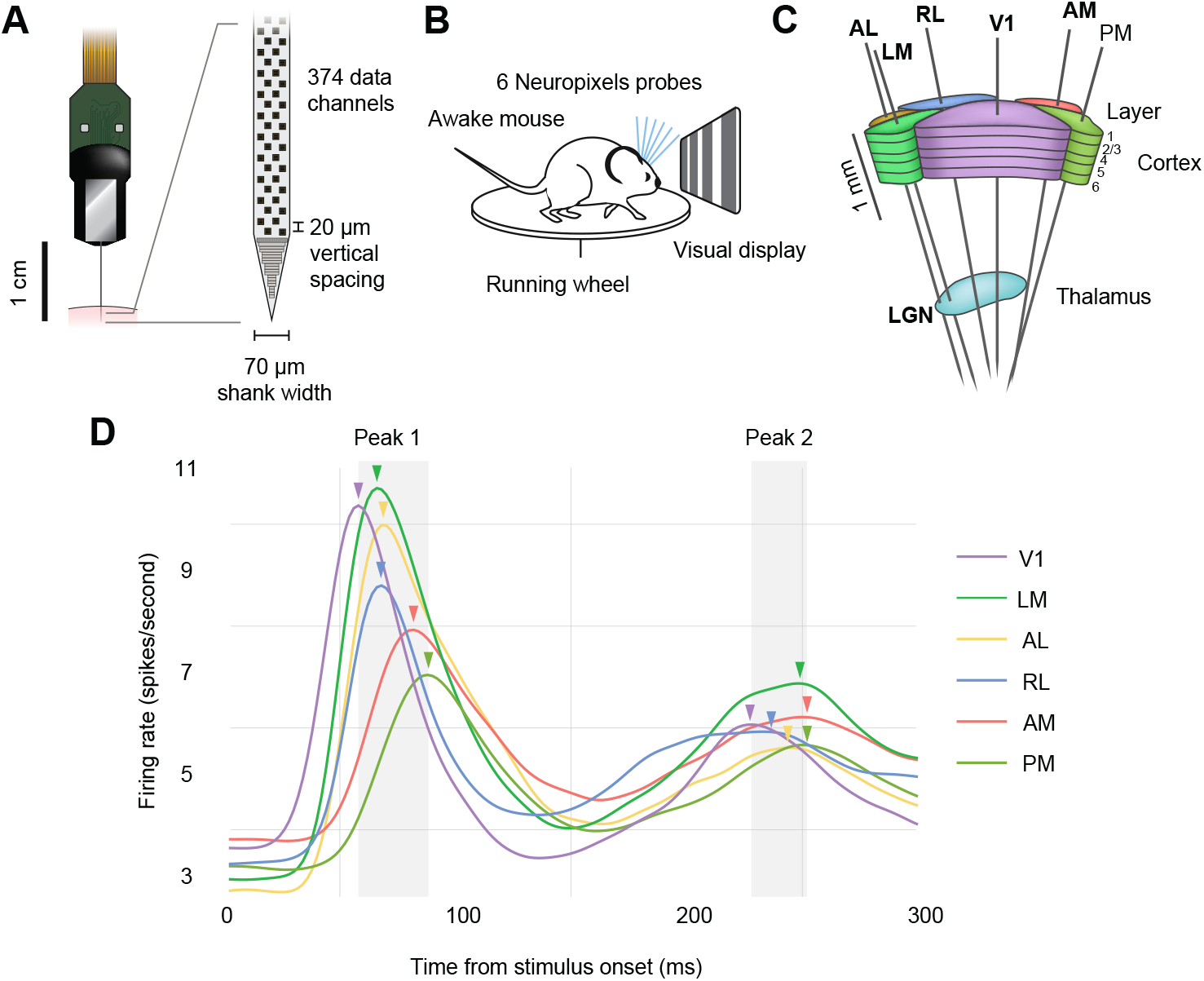
Electrophysiological recordings from seven visual areas in a publicly available dataset. **A**, Illustration of a Neuropixels probe used to detect extracellular spiking activity across hundreds of neurons in parallel. **B**, Schematic of the recording configuration. Mice are head-fixed and free to run on a spinning wheel, while passively exposed to visual stimuli. Six Neuropixels probes are targeted to the visual cortex. **C**, In each recording session, probes pass through six visual cortical regions (AL, anterolateral visual area; LM, lateromedial visual area; RL, rostrolateral visual area; V1, primary visual cortex; AM, anteromedial visual area; PM, posteromedial visual area) and one thalamic visual region (LGN, lateral geniculate nucleus). **D**, Overall population response to the onset of a drifting grating stimulus. The population response here is obtained by smoothing PSTHs across neurons and trials for each area. Note the two prominent peaks, which likely result from feedforward and feedback signal propagation, respectively. Arrows indicate the time at (and thus order in) which the firing rate in each area’s population reaches its maximal value.

We followed Chen et al. (2022) by focusing on the time at which each population firing rate reaches its maximum value (“peak timing”) because, by definition, many spikes occur around the time of the maximal value, so it should be well determined, statistically. Chen et al. Chen et al. (2022) noticed that the simple (“näive”) method of determining peak timing, namely smoothing the population PSTH and finding the time of its maximal value, in fact, produced poor estimates, and they identified three sources of difficulty. First, for any given condition, only a subset of neurons responded similarly to produce the two-peak population profile, while the other neurons effectively diluted the signal by issuing irrelevant noisy spike times. Second, the shape of the peak was condition-specific; the usual population PSTH, as shown in Figure 1, is a blurred average across conditions. Third, the Poisson-like noise in the spike trains (which is typical in many brain areas of behaving mammals), from a limited number of condition-relevant neurons, contributed substantially to the inaccuracy in peak timing estimates derived from smoothed population PSTHs. We devised an analytical strategy to overcome these problems, and the new method is much simpler and more computationally efficient than that proposed by Chen *et al*.

Not only do excessively noisy estimates of timing make it difficult to establish sequential activity across areas, they can also greatly decrease correlations of the timing across two areas, such as V1 and LM. This phenomenon is well known and easy to prove mathematically (Kass et al., 2014, Section 12.4.4). It is also intuitive: if two measurements tend to move up and down together but independent noise is added to them, the extent to which they move together will be thrown off by the noise, and their correlation will thus be diminished. In the statistics literature, an improved correlation estimate (often discussed under the heading of “errors in variables”), is typically called a “correction for attenuation” (Kass et al., 2014, Section 12,4.4 and references therein). To be clear, attenuation of correlation results from sampling a relatively small number of neurons: our corrections for attenuation aim to do a better job of estimating the results that would have been obtained had the entire condition-relevant population of neurons been recorded. Figure 6 provides an illustration for areas V1 and LM, based on the method we describe here.

Chen *et al*. solved the three problems listed above by developing a comprehensive Bayesian hierarchical model, called the Interacting Population Rate Function (IPFR) model. Simulation studies showed their method could obtain accurate estimates of individual trial population burst times and their trial-to-trial correlations across areas. Because it included, together, all elements of the problem, the IPFR model was rather complicated, and for large data sets could take an excessively long time to run in standard computing environments. As an alternative, we developed a simplified version by solving each of the three problems, separately, in a 3-step procedure. We demonstrate that the new procedure can replicate, with good accuracy, the results of the previous method while having an 85 to 90% reduction in compute time. We then use the new procedure to examine the relative timing and coupling of population bursts across thirteen mice and to infer the mouse-to-mouse variation in these timing and coupling relationships.

## Materials and Methods

### Experimental Setup

In this work, we analyzed the publicly available Allen Brain Observatory Visual Coding Neuropixels Dataset Allen Institute MindScope Program (2019), which includes spike trains and local field potential recordings from the mouse visual system. In each experiment, six Neuropixels probes were targeted to six areas of visual cortex (Figure 1A-C), which were identified via functional retinotopic mapping before the experiment. Spike trains from between 40 and 100 neurons from each area were recorded simultaneously from each subject (after applying standard thresholds to spiking sorting quality metrics, see Siegle et al. (2021) for details). Thirty mice were head-fixed and passively presented with visual stimuli, which included natural movies, full-field flashes, Gabor patches, and drifting gratings. Here, we focused on drifting gratings because they included many repeated trials for each condition, the trials are relatively long (3 s each), and they drive strong responses in visual cortex. The drifting gratings have 40 conditions that result from combining eight grating orientations (0°, 45°, 90°, 135°,180°, 225°, 270°, 315°) and five temporal frequencies (1, 2, 4, 8, 15 Hz). Each condition is repeated 15 times. Each trial lasts for 3 s, with 2 s stimulus and 1 s blank screen, with all conditions randomly interleaved. We analyzed spike trains from the lateral geniculate nucleus (LGN), the thalamic region that receives inputs from the retina and sends outputs to cortex, and six cortical areas: primary visual cortex (V1), which is the primary target of LGN, and is at the bottom of the visual hierarchy, as well as the rostrolateral (RL); lateromedial (LM); anterolateral (AL); anteromedial (AM); and posteromedial (PM) visual areas, with the last two residing at the top of the anatomically defined visual hierarchy Harris et al. (2019); D’Souza et al. (2022).

### Model Overview and statistical analysis

A high-level sketch of the IPFR model for a single area, under a single stimulus condition, is shown in the left diagrams of Figures 2 and 3, and details can be found in Chen et al. (2022). The three steps of our new procedure correspond to the three problems identified in the introduction. We label these steps (1) interacting population selection, (2) initial peak time and standard error estimation, and (3) peak time denoising using and trial-to-trial correlation. Condition-specific population selection produces better timing estimates by removing irrelevant spikes produced by neurons with low activity, but it leaves us with comparatively small samples of good neurons. The denoising (in step 3 based on input from step 2) is needed to compensate for the noisiness of small-sample estimates. In the abstract ideal where an entire relevant population would be recorded, the denoising would not be necessary. In real-world experiments, however, noise obscures correlations. Schematic summaries of this procedure are shown in Figures 2 and 3.

**Figure 2:**
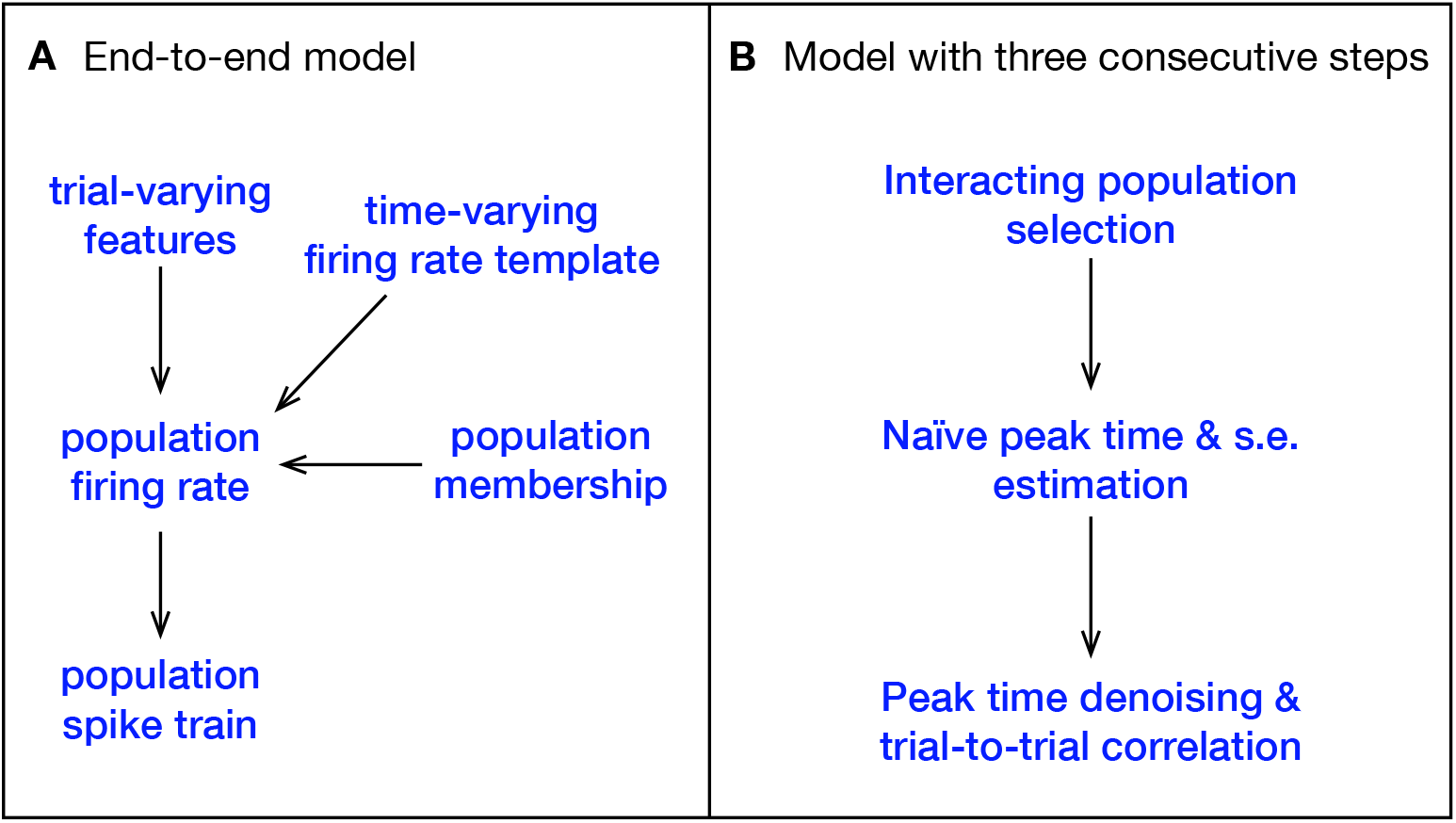
Comparison between the IPFR model and our three stage model for a single stimulus condition. **A**, The IPFR model. The population spike train on a single trial is driven by its population firing rate, which combines a time-varying firing-rate template with trial-varying features. Only a subset of neurons recorded within the brain area will be used, and this subpopulation is determined by a population membership probability. This is all captured by a single model, with all variables and parameters jointly inferred. **B**, In our model, the estimation procedure is divided into three sequential stages.

**Figure 3:**
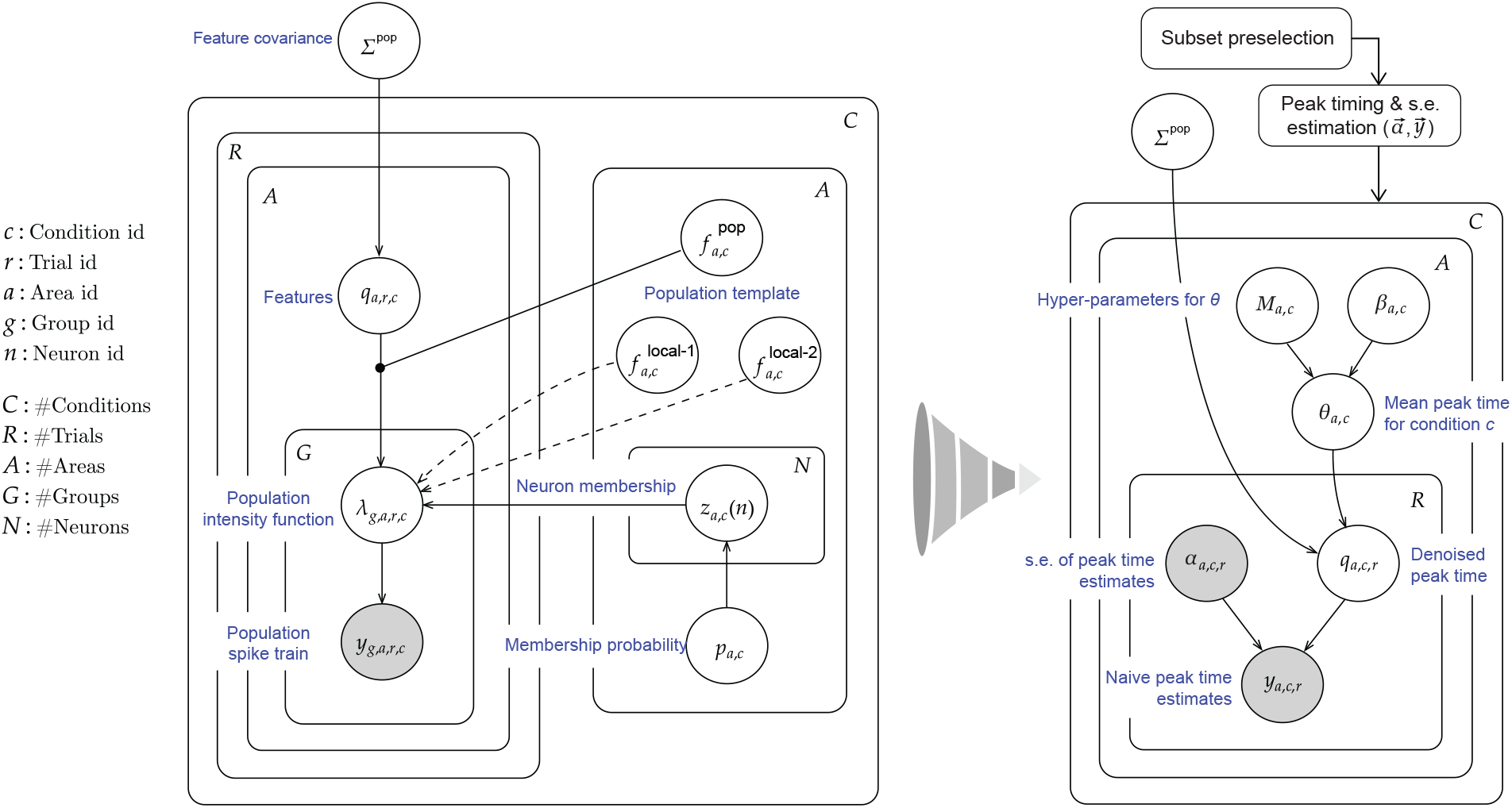
Plate diagram of our simplified multi-step procedure for estimating the timing of population bursts. Left: Original IPFR model from Chen et al. (2022). Right: Simplified model, which divides the estimation task into 3 steps. We are able to obtain comparable results with substantially reduced computation time.

In step (1) we extracted the subset of the population in each area that responds to a particular condition, which we call the interacting population. As we were interested in the time from stimulus onset at which the intensity function reaches its peak, we filtered out neurons that showed no change in firing rate, or a decrease in firing rate, in response to the stimulus. Then we selected, for each condition, the neurons with a clear peak in the stimulus response profile. We accomplished this selection by fitting a firing rate function to the PSTH for each recorded neuron, and for each stimulus condition, across trials. A neuron’s condition-specific PSTH was obtained by first binning the spike train in each trial into 1 ms bins, and then summing these binned spikes trains across all the condition’s trials. The firing rate function is modeled non-parametrically using a Poisson Generalized Additive Model (GAM) with a spline basis (Figure 4), which is fit to the neuron’s PSTH using maximum likelihood (Kass et al., 2014, Chapter 19).

**Figure 4:**
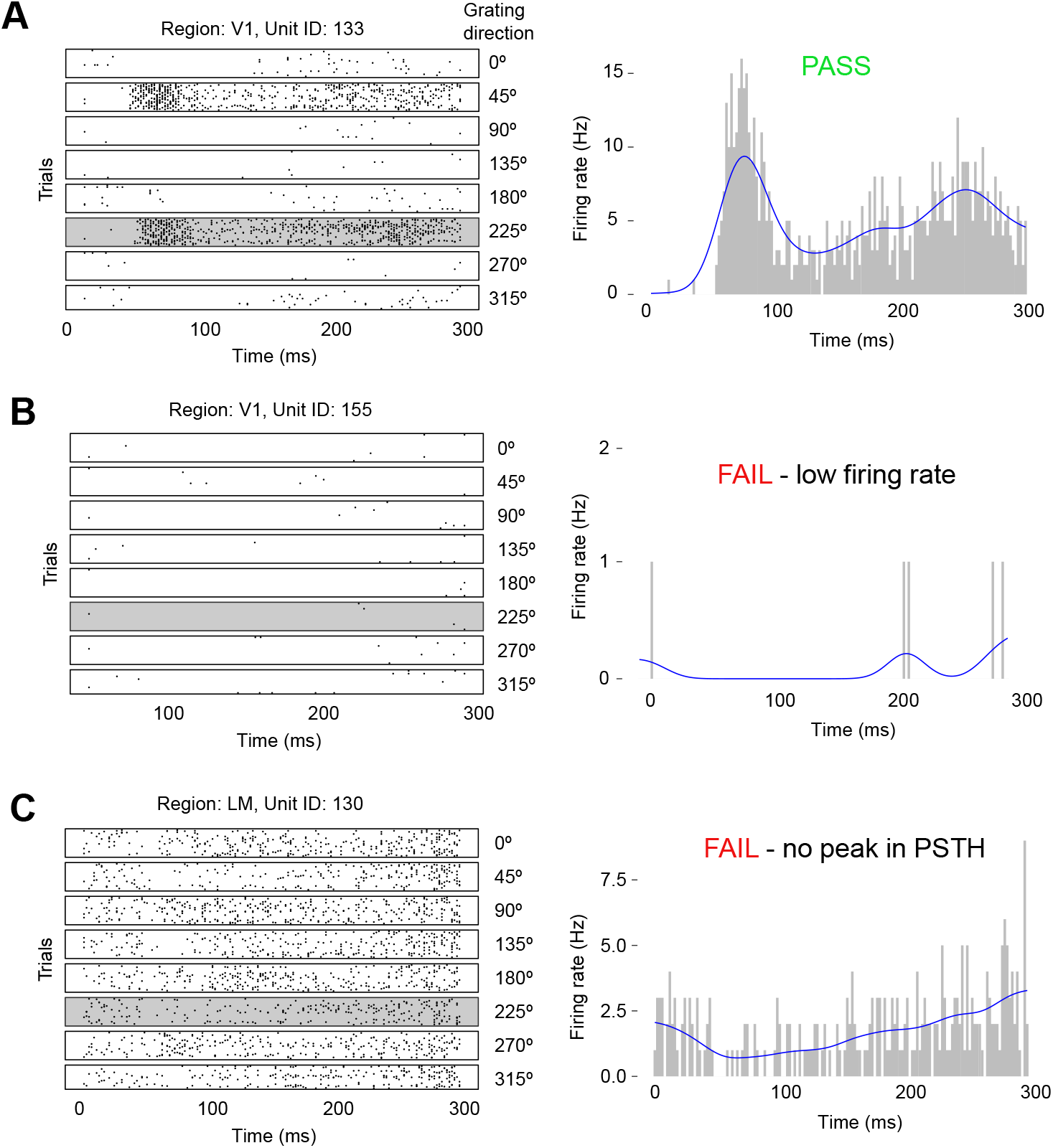
Selection criteria illustrated for three example neurons. Left: Spike rasters of three different neurons to eight directions of the 1 Hz drifting grating stimulus. Right: PSTH and fitted firing rate function for the 225 degree stimulus condition. **A**, The neuron passes the selection criteria due to its high firing rate and peak in its stimulus response profile. **B**, The neuron fails the selection criteria due to its low firing rate. **C**, This neuron fails because its PSTH lacks a peak (defined as a concave critical point).

We observed strong orientation tuning in the response profiles of individual neurons, consistent with previous recordings from visual cortex. Examples are shown for three neurons in Figure 4. Exploratory analysis revealed that the initial population burst occurs between 30 ms and 160 ms from stimulus onset, and so filtering for the evoked responses typical of an interacting population is done within this window. The filtering criteria for each neuron’s firing rate function within the burst window are as follows:

(i) average firing rate is in the top 60% among neurons in the same visual area;
(ii) must have a concave critical point;
(iii) must have maximum slope in the top 60% among neurons in the same visual area; and
(iv) the increase from baseline to peak firing rate in the top 60% among neurons in the same visual area.

These filtering criteria can be easily automated and applied to all mice. Intuitively, they select the neurons within each visual area with a strong, peaked response to a stimulus. The conditions for a strong peaked response were determined from exploratory analysis on a single, randomly selected mouse, to be a high average firing rate (across time) in the peak response time window, a concave peak, and a sharp and noticeable increase in firing rate from its baseline in the peak response window. The presence of these features together is a strong indicator of a peaked response, although the concavity and the increase from baseline matter to a greater extent than the absolute firing rate. The thresholds were determined empirically for each peak separately from an exploratory analysis of data from one mouse (ID 756029989) and validated on a second mouse (ID 760345702) before extending the analysis to the full Allen dataset.

After filtering, we rejected data from any condition in each of the 7 visual areas with an interacting population of less than 10 neurons available for the next stage of the analysis. Following the subset pre-selection step, the resulting data set for each visual area consists of only those neurons that contribute to population activity given each stimulus condition.

In step (2), the population PSTH on a given trial for a particular area is obtained by summing binned spikes trains across neurons. The population firing rate function is estimated in the same manner as with individual neurons (using a Poisson Generalized Additive Model with a spline basis), and the time of maximal firing rate, which is a näive estimate of the “peak time”, is obtained from the population firing rate function as the time at which the maximum of this function occurs; its estimation uncertainty is obtained by bootstrap resampling from the population of neurons. As an object of statistical estimation, the peak burst time has the advantage of having, by definition, a relatively large number of spikes occurring near that time.

In step (3), the naive peak times and uncertainties obtained in step (2), which are represented as *y*_*c,r*_ and 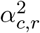 respectively, are inserted into a simple Bayesian hierarchical model, shown visually in figure 3 and specified as follows, for trial *r* under condition *c*:

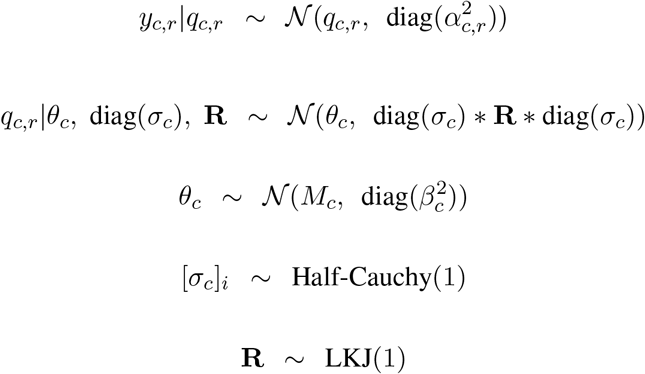

The components of the vectors correspond to the visual areas, indexed by *a* in figure 3. Here, *y*_*c,r*_ is a vector of peak time estimates (found in step (2)), which are assumed independent and identically distributed given *q*_*c,r*_. The components of the vector 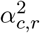 (again found in step (2)) are squared standard errors of the peak time estimates. The components of the vector *q*_*c,r*_ are denoised peak times (expected values of the components of *y*_*c,r*_, which may be considered the vector of peak times based on the hypothetical complete populations of condition-relevant neurons) while *θ*_*c*_ are mean denoised peak times (the expected values across trials) and *σ*_*c*_ are the corresponding standard deviations, for condition *c*.

To complete the hierarchical model we adopt prior probability distributions that are commonly used because of their good statistical behavior. We place a half-Cauchy distribution on the standard deviation, with scale parameter 1 Polson and Scott (2012); Gelman (2006). The symbol **R** denotes the matrix of cross-area correlations in the peak-1 times. For its prior we use a probability distribution over the space of (flattened) vectors of product-moment correlations that composes **R**. This is called the Lewandowski-Kurowicka-Joe (LKJ) distribution Lewandowski et al. (2009), and is the recommended prior distribution over correlation matrices in the popular Bayesian inference software package STAN Stan Development Team (2024). This distribution has constant density over the component representation of the manifold of *d*-dimensional positive definite symmetric matrices with unit diagonals and off-diagonals between -1 and 1. Note that this manifold has a non-Euclidean geometry because of the constraints imposed by symmetry, unit diagonal, and positive definiteness. A sampling algorithm proposed in Lewandowski et al. (2009) can generate samples from this manifold, and therefore defines a probability density function over the manifold. The density for a correlation matrix sampled according to this algorithm is proportional to the determinant of the matrix raised to a certain power, which is defined by a free parameter. When this power is 0, the probability density corresponds to a uniform distribution over the component representation of this manifold. The matrix 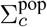 in the right-hand side of Figure 3 can be expressed as diag(*σ*_*c*_) * **R** * diag(*σ*_*c*_), and is the cross area covariance matrix for the peak times *q*_*c,r*_. The symbols *M*_*c*_ and *β*_*c*_ are the mean and variance hyper-parameters for the prior on *θ*_*c*_. We estimate both using maximum likelihood, in a similar way as *y*_*c,r*_ and *α*_*c,r*_, by summing binned spike trains across both neurons and trials corresponding to stimulus condition c, for each area. This closely approximates fully Bayesian posterior inference, as described in the conditionally independent hierarchical model (CIHM) framework Kass and Steffey (1989), also referred to as parametric empirical Bayes (PEB). We use the Rstan package with Hamiltonian Monte Carlo Stan Development Team (2024) to obtain posterior samples and then posterior means and variances for *q*_*c,r*_, *θ*_*c*_, *σ*_*c*_ and **R**, *r* = 1, …, *R, c* = 1, …, *C*. In addition, we use the posterior mean peak times 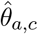 and posterior variances 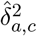 to compute, for each area *a*, estimates of the mean peak time across conditions, 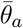. We do this using a weighted mean as the differing degrees of variances in the peak time estimates across areas makes the simple arithmetic mean a higher variance estimator. We do not use the usual formula for a weighted mean (e.g., page 193 of Kass et al. (2014)) because 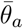 involves two sources of variance, the posterior variances 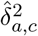 of the condition-dependent estimates and the variance of those estimates across conditions. Thus, the formula we need appears in equation (16.37) on page 461 of Kass et al. (2014). We use the following simple iterative algorithm to estimate the appropriate weighted mean and its variance (the maximum likelihood estimate of these two quantities) by alternating between the two, while conditioning on the current value of the other:

- Initialize

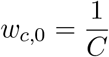
- For k = 1 till convergence, repeat

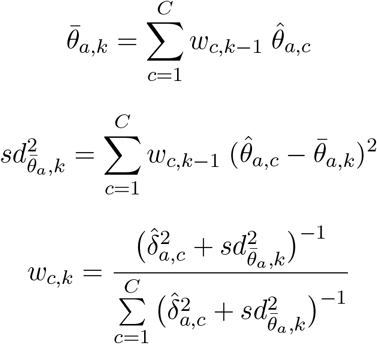

where 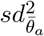 is an estimate of the variance in the peak times of area *a*, across stimulus conditions *c* = 1, …, *C*, around their weighted mean 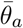.

We compute the squared standard error of the weighted mean, 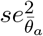 as

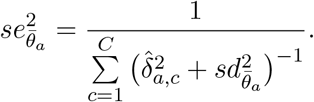

We follow these steps for each mouse to get the weighted means and variances, for all areas. Based on these, for each area, we then aggregate across all mice using the ordinary weighted mean and its standard error; we also compute the standard deviation across mice. We do this analysis for the first and second peaks separately.

## Data and code availability

The code for our model is available [on GitHub](https://github.com/Tolani-O/Fast_timing_and_coupling_estimation). The data from the Allen Brain Observatory Neuropixels Visual Coding dataset can be accessed via the [AllenSDK](https://allensdk.readthedocs.io/en/latest/visual_coding_neuropixels.html), the [DANDI Archive](https://dandiarchive.org/dandiset/000021), and through [AWS Registry of Open Data](https://registry.opendata.aws/allen-brain-observatory/).

## Results

In order to establish the estimation accuracy of our model, we first conducted simulations using data generated by the IPFR model. We demonstrate that our three-step method is nearly as accurate as the IPFR model while having greatly reduced computation time.

We then apply our method to data from the Allen Brain Observatory Neuropixels Visual Coding dataset, with the goal of examining mouse-to-mouse variation in relative lead–lag timing and coupling (trial-to-trial correlation) relationships among the different visual areas based on peak 1 and peak 2 timing. Previous studies have shown a temporal ordering in the feedforward propagation of spikes through the visual cortex, with evoked spikes appearing in higher visual areas having longer delays after stimulus onset Schmolesky et al. (1998); Siegle et al. (2021); Glickfeld and Olsen (2017). We expected such an ordering to be reflected in the ordering of peak 1 times across the different areas. Furthermore, any ordering that is functionally relevant should be consistent across subjects, even though the absolute peak times may be subject to a variety of sources of variation having little or no functional relevance. Because feedback propagation has been less well studied, it is unclear what to have expected about the ordering of peak 2 population activity, as peak 2 timing most likely depends on top-down signals coming from other brain regions, as well as the animal’s internal state.

In addition to the relative ordering in the peak times across the visual areas, our methods can be used to learn about functional associations between visual areas through marginal and partial trial-to-trial correlations in peak times between pairs of areas. For example, neuroanatomical studies of the mouse thalamocortical pathway have shown there are almost no direct anatomical projections from LGN to higher visual areas Antonini et al. (1999), which implies communication between the LGN and the higher-order visual areas is mediated through V1. We would therefore expect any marginal correlations between LGN and higher-order areas to be reduced substantially after accounting for the activity of V1, using partial correlation. We report results using these tools.

### Performance in estimating ground truth values using simulated data

To compare our proposed model to the IPFR model, we conducted a simulation study using the latter model as the ground truth. Our simulation consisted of two hypothetical brain regions *a*_1_ and *a*_2_, with the number of neurons in each area given by *N*_1_ and *N*_2_ areas, respectively. We specified a stimulus condition *s*, with proportions 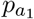 and 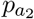 of the neurons in the corresponding region belonging to the single peaked response population for *s*, and the complementary sets of neurons in each region belonging to the flat response population. Given these conditions, as well as the pre-defined peaked and flat response population firing rate templates for each region, we sampled the individual neuron spike trains using a Poisson point process. For a given area, each neuron spike train depends on the neuron’s population membership (which follows a categorical distribution), and corresponding population firing rate template. In addition, each trial of the peaked response population firing activity depends on a trial-varying peak time, which follows a bivariate Gaussian distribution (corresponding to each the two regions), with a pre-specified mean *μ*, variances *σ*_1_ and *σ*_2_ and correlation *ρ*. We incorporated the trial peak time into the peaked response population firing rate function using time warping. Details of the generative model can be found in Chen et al. (2022). We sampled the neuron spike trains in both areas for *R* trials, and then fit the IPFR model and our three-step model to the resulting data. We also estimated peak times from a Gaussian kernel applied to the population PSTH (we referred to this previously as a näive estimate of the peak times), with bandwidth selected using cross-validation. The Gaussian kernel, also called Gaussian filter, is commonly used in practice to estimate a firing rate function. We therefore consider it a baseline approach. We repeated this process of simulating and fitting data across 60 repititions, for each of several configurations of *ρ, R* and lag (= *μ*_2_ −*μ*_1_). We fixed *N*_1_, *N*_2_, *σ*_1_, *σ*_2_, 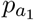, 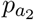 at the values 100, 100, 1, 1, 0.8, 0.8, respectively. For each repetition we estimated the parameters of the Gaussian distribution of peak times using each method (for the näive method we estimated correlations from the Pearson correlations of the näive peak time estimates), and then computed the mean and standard error across repetitions. Figure 5 shows the fitted firing rate function for each candidate model to area *a*_1_’s population PSTH in a single trial of an example simulated dataset, shown for visual comparison.

**Figure 5:**
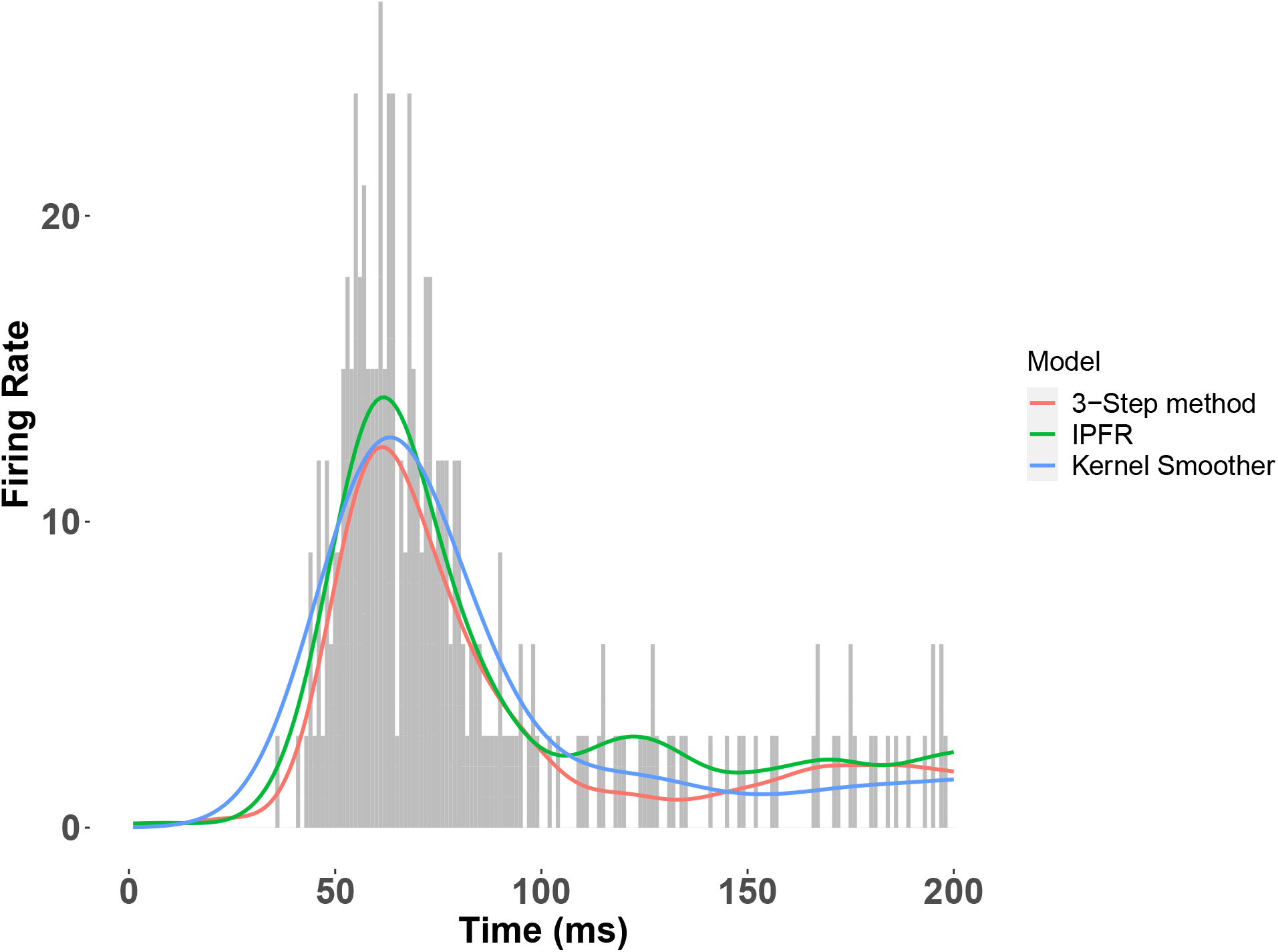
Fitted firing rate function for each candidate model to the population PSTH in a single trial of an example simulated dataset. For one of the datasets with correlation *ρ* = 0.8 and number of neurons *N*_1_ = 100, we show the fitted firing rate function on a single trial, using each method described above. The IPFR and the 3-step method both use a GAM with a log link function to fit the intensity function. However, the 3-step method first filters out those neurons that do not participate in the population burst response, as detailed in the Model Overview and Statistical Analysis section above. Note that the IPFR (green trace) was used as the ground truth to generate the datasets.

The IPFR and the 3-step method both use a GAM with a log link function to fit the intensity function. However, the 3-step method first filters out those neurons that do not participate in the population burst response, as detailed in the Model Overview and Statistical Analysis section above. Tables 1 and 2 show the outputs, summarized across 60 simulated datasets, for each candidate model. In Table 1, the lag times estimates produced by the kernel smoother are close to those from the other two methods. However, the standard error bars are about twice as large as those obtained from the 3 step method. Thus, using the 3-step method would correspond to results obtained from the kernel smoother based on four times as much data. Taking the kernel smoother as the reference, Table 2 shows the percentage reduction in estimation error obtained from each method when estimating trial-to-trial correlation. While our method does not have as much of an improvement over the kernel method as the IPFR model on data simulated from the IPFR model, it still considerably improves over the reference, demonstrating the ability of our method to denoise the trial peak times and to more accurately estimate the correlations.

**Table 1:** Lag recovery (in milliseconds) from three methods from data simulated from the IPFR model. In two hypothetical brain areas, and for one stimulus condition, we simulated neuron spiking data, using the IPFR model as the ground truth, for different average lag in peak time between the two areas. We kept *ρ, R, σ*_1_, *σ*_2_, *N*_1_, *N*_2_, 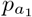 and 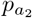 fixed at 0.8, 60, 1, 1, 100, 100, 0.8 and 0.8 respectively. We applied the three methods to recover the ground truth lags, and computed the mean estimate for each method across 60 repetitions, as well as the simulation standard errors, shown in parenthesis. We note that in our simulated datasets, the kernel smoother itself produces a decent estimate of the lag time, but has standard errors that are twice as large as the other two methods.

| Parameters |  | Model avg. estimate (s.e) |  |  |
| --- | --- | --- | --- | --- |
| lag |  | IPFR model | 3-step method | Kern smooth |
| 8 |  | 8 (0.08) | 8 (0.1) | 7.9 (0.26) |
| 0 |  | 0 (0.07) | 0 (0.11) | 0.02 (0.21) |

**Table 2:** Correlation recovery from three methods from data simulated from the IPFR model. Same procedure as in Table 1, but here we used different combinations of trial-to-trial correlations *ρ*, and number of trials *R*, with lag, *σ*_1_, *σ*_2_, *N*_1_, *N*_2_, 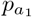 and 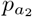 held fixed at 8ms, 1, 1, 100, 100, 0.8 and 0.8 respectively. We applied our three step method, the IPFR model, and a naïve model based on kernel-smoothed population intensities, to 60 simulated datasets, to recover the ground truth correlations. We compute the mean estimate for each method across repetitions and the simulation standard errors, shown in parenthesis. We also computed the percentage reduction in the estimation error from the naïve kernel smoother achieved by both ours and the IPFR model.

| Parameters |  | Model avg. estimate (s.e) |  |  | % error reduction |  |
| --- | --- | --- | --- | --- | --- | --- |
| $\rho$ | R | IPFR model | 3-step method | Kern smooth | IPFR model | 3-step method |
| 0.8 | 60 | 0.79 (0.05) | 0.78 (0.08) | 0.32 (0.11) | 98 | 96 |
| 0.2 | 60 | 0.19 (0.1) | 0.18 (0.12) | 0.08 (0.19) | 92 | 83 |
| 0.8 | 30 | 0.79 (0.07) | 0.75 (0.16) | 0.27 (0.19) | 98 | 91 |

In order to demonstrate the improvement in runtime between the IPFR and the 3-step method, we ran additional simulation studies on the previously described models. First, we report the difference in runtime of both models for number of simulated areas = 2, 4, 6, and 8. For each value, we specify *N*_*i*_ = 100 neurons in each area. The results are summarized in Table 3. Increasing up the number of areas also increases up the number of neurons to be assigned to a peaked or flat response population, as well as the size of the covariance matrix being estimated. We used a randomly chosen mean vector and covariance matrix for the trial peak time distribution, and we used a peaked response population proportion of *p*_*i*_ = 0.8 for each area. From the results in Table 3, we observe between 88 and 90% reduction in the runtime of the 3-step method over the IPFR. We also separately investigate the relative runtimes of both models as we vary the number of stimulus conditions, which also varies the number of trials. Table 4 summarizes our results for number of stimulus conditions = 1, 5 and 10. Fixing the number of areas at 2, and the number of neurons per area at 100, we use *R*_*s*_ = 60 trials for each stimulus condition *s*_*k*_. Under this configuration, the population membership of the *jth* neuron in area *a*_*i*_ now also depends on the stimulus condition *s*_*k*_ being considered. We set the proportion of peaked response neurons *p*_*i,s*_ = 0.8 for each area *a*_*i*_ and stimulus conditions *s*_*k*_, and the peak time trial to trial correlation and variance as *ρ* = 0.8 and *σ* = 1 respectively. In both scaling experiments, we generated 4000 samples of each variable of interest. While both models scale quadratically with both the number of areas and stimulus conditions (number of neurons and trials, respectively), we obtained between an 85 and 90% reduction in runtime with the 3-step method over the IPFR for problems of the same size. We note that the simulated datasets used in this study are quite small (a single stimulus condition in the area scaling experiment, two areas in the stimulus condition scaling experiment) compared to interesting real-world datasets (5 to 10 areas and 10s of stimulus conditions on multiple mice), and thus the model time complexity becomes progressively more important on the scales of real-world data. Extrapolating these results to the 7 areas and 40 stimulus conditions used for each mouse in this study, fitting the model would have taken more than 18 hours with the IPFR method, versus 3 hours with our 3-step method.

**Table 3:** Runtime comparison between the IPFR and the 3-step method with a varying number of areas. We simulated neuron spiking data, using the IPFR model as the ground truth, for A = 2, 4, 6, and 8 areas, each with 100 neurons. We had s = 1 stimulus condition, with R = 60 trials. We used a randomly chosen mean vector and covariance matrix for the Gaussian trial peak time distribution. The proportion of neurons in each area belonging to the peaked response population was fixed at 0.8. We applied the IPFR and the 3-step method to 10 simulated datasets, running the algorithm for 4000 iterations in each instance. We measured the average runtime in both models, as well as their standard errors across repetitions, shown in parenthesis. We also computed the percentage reduction in runtime obtained by the 3-step method over the IPFR.

| Model avg. runtime in mins (s.e) |  |  |  |
| --- | --- | --- | --- |
| # Areas | IPFR model | 3-step method | % runtime reduction |
| 2 | 192 (11.3) | 23 (4.3) | 88 |
| 4 | 278 (14.7) | 31 (3.9) | 89 |
| 6 | 349 (12.3) | 35 (4.7) | 90 |
| 8 | 406 (15.8) | 44 (3.6) | 88 |

**Table 4:** Runtime comparison between the IPFR and the 3-step method with a varying number of stimulus conditions. Similarly to table 3, we simulated neuron spiking data, using the IPFR model as the ground truth, for s = 1, 5 and 10 stimulus conditions, each with 60 trials, for A = 2 areas with a 100 neurons in each. The trial-to-trial correlations *ρ*, lag, *σ*_1_ and *σ*_2_ were held fixed at 0.8, 8ms, 1 and 1 respectively. The proportion of neurons in each area belonging to the peaked response population was fixed at 0.8 for all stimulus conditions. We applied the IPFR and the 3-step method to 10 simulated datasets, running the algorithm for 4000 iterations in each instance. We measured the average runtime in both models, as well as their standard errors across repetitions, shown in parenthesis. We also computed the percentage reduction in runtime obtained by the 3-step method over the IPFR.

| Model avg. runtime in mins (s.e) |  |  |  |
| --- | --- | --- | --- |
| # Stimulus conditions | IPFR model | 3-step method | % runtime reduction |
| 1 | 184 (10.3) | 21 (5.3) | 89 |
| 5 | 221 (10.8) | 31 (5.9) | 86 |
| 10 | 307 (9.2) | 45 (6.2) | 85 |

### Illustration with real data

We chose one example subject, and two visual areas V1 and LM, to demonstrate the method’s correction for attenuation of correlation, and the extent to which this is facilitated by both the sub-population selection and the denoising. Results are shown in Figure 6. We estimated the Pearson correlation coefficient for the trial-by-trial peak 1 times in three cases. First (panel A), peak times were obtained by applying a kernel smoother on each trial to the PSTH based on the full population (without the interacting population selection step). The correlation was .06. Second (panel B), we applied the kernel smoother to the PSTH after selecting the interacting population. The correlation increased to 0.2. Third (panel C), the peak times were obtained as posterior means from the Bayesian hierarchical model, according to the complete three-step approach. The correlation obtained further increased substantially to 0.8. The results from our simulation studies suggest that the correlation value of 0.8 were likely attenuated to 0.06 in the näive estimates shown in panel A. See Behseta et al. (2009) for further depictions of attenuation of correlation in multi-trial spike count data. This example case illustrates the importance of the denoising step in our procedure.

**Figure 6:**
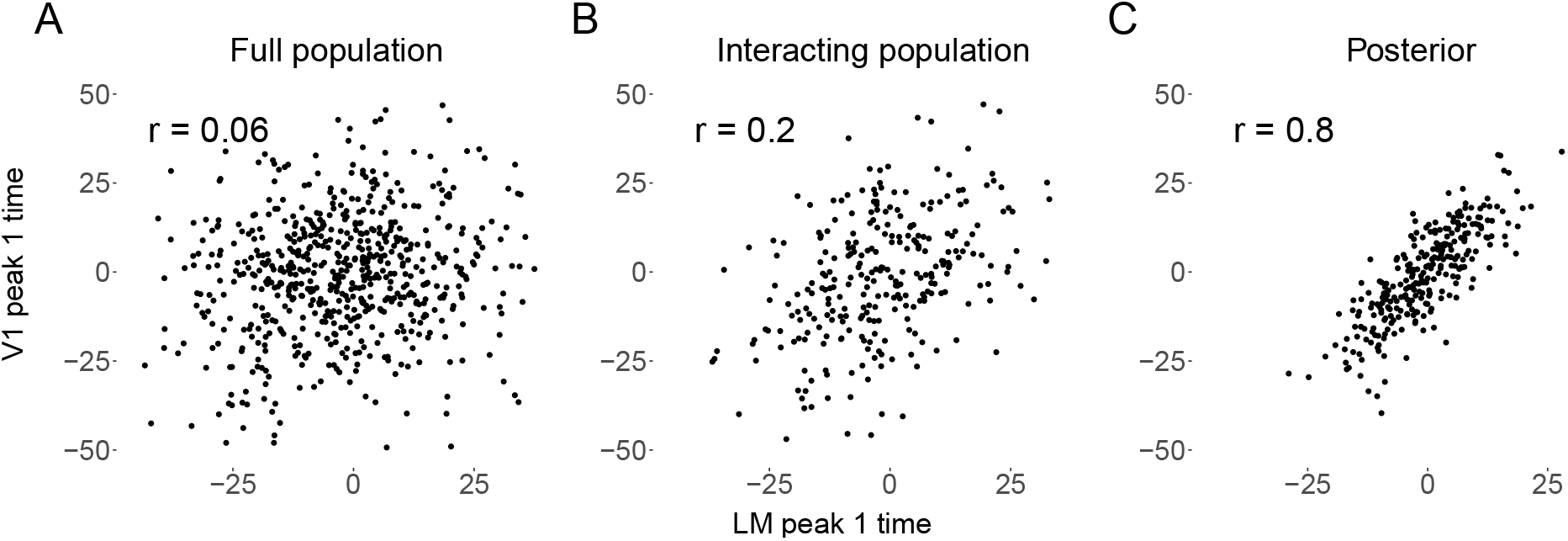
Denoising of peak 1 times for regions V1 and LM in an example mouse. **A:** Plot of Estimated peak 1 times using a kernel smoother applied to the condition-specific PSTH based on the full populations of recorded neurons. **B:** Plot of estimated peak 1 times after interacting sub-population selection. **C:** Plot of estimated peak 1 times after applying the full three-step method.

### Analysis of data from multiple mice

To understand the functional ordering present in feedforward signal propagation in the visual cortex, we applied our method to data from thirteen mice from the Allen dataset, estimating the average trial-by-trial peak times in each of seven areas. We then computed the standard deviations of the peak time across mice for each area. Figure 7 shows the weighted means, standard errors, and standard deviations for peak (1 and 2) times across mice. While the ordering of Peak 1 times in Figure 7 is consistent with previous results (see Figure 3 of Siegle et al. (2021)), that figure can not indicate the extent to which such timing is or is not consistent across subjects. We examine consistency next.

**Figure 7:**
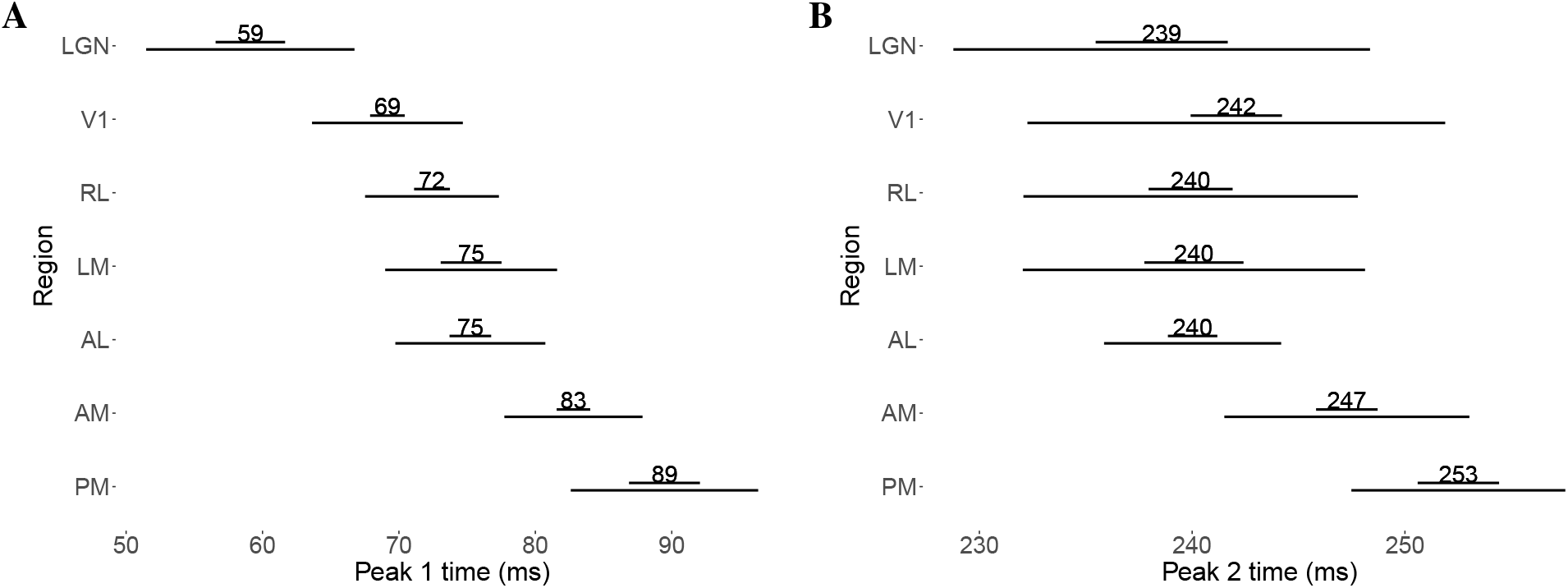
Weighted means, standard errors, and standard deviations across mice for peak 1 time and peak 2 time. The shorter horizontal bar (top) represents the standard errors, and the longer bars (bottom) represent the standard deviations, **panel A** peak 1 (13 mice), **panel B** peak 2 (11 mice). The ordering in peak 1 times largely disappears in peak 2 times, except that areas AM and PM appear to have somewhat later peak 2 times.

#### Consistency in ordering of peak times

Despite subject-to-subject variability in actual peak time values, Figure 8 shows some consistency in the ordering of peak times across mice. The figure also displays some inconsistencies. For peak 1, across all mice, we observe LGN preceding V1, which precedes higher-order visual areas. For all mice we also observe peak 1 time in both AM and RL preceding PM. However, apart from these relationships, the relative peak 1 timing among higher-order visual areas is specific to each mouse. For peak 2, for all mice LGN and V1 both precede AM and PM, and again RL precedes PM, but all other orderings are inconsistent. For example, LGN precedes V1 sometimes, but not uniformly across mice, and V1 precedes higher-order areas other than AM and PM sometimes, but not for all mice. Again, as with Peak 1, the relative timing among higher-order areas apart from RL and PM is inconsistent.

**Figure 8:**
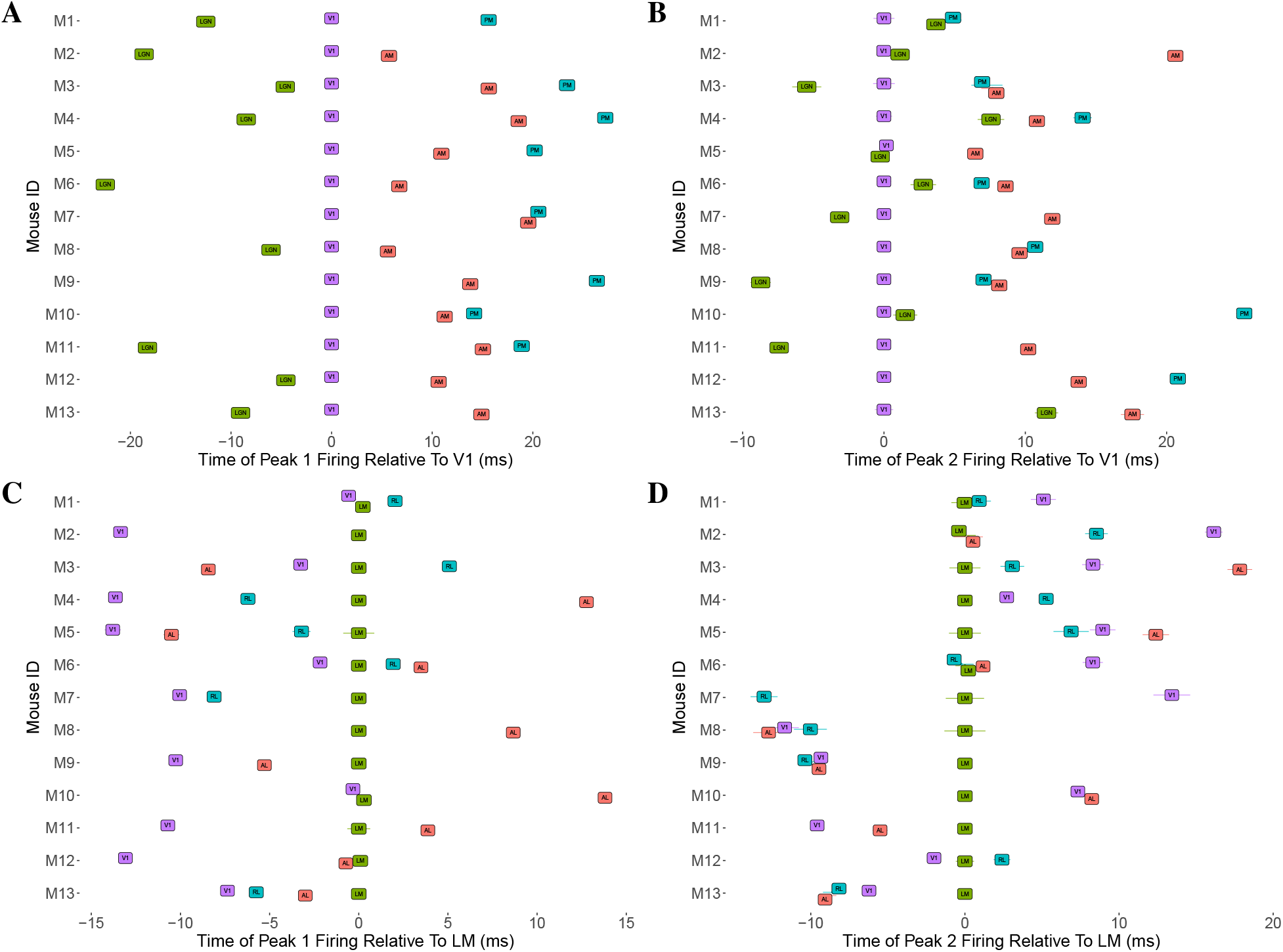
Mouse-to-mouse variability in the time of the initial peak response relative to a reference region. In panels **A** and **B**, we show Peak 1 and peak 2 (respectively) time estimates for LGN, AM and PM, relative to the corresponding peak time estimate for V1, for the same set of thirteen mice. Panels **C** and **D** show the peak 1 and peak 2 times estimates for V1, RL, LM and AL, relative to the corresponding peak time estimate for LM. In all cases, 1 standard error bar is also shown for the peak time estimates, although many are small enough to be obscured by the region label. We observe a consistent ordering across mice in the peak 1 times of LGN, V1, AM and PM in **A**, suggesting a functionally relevant pathway. We don’t observe the same consistency in the peak 2 times for these areas in **B**, although we see LGN and V1 consistently reach their second peak before AM and PM. Among the regions V1, RL, LM and AL, we see that for peak 1 in **C**, V1 tends to reach its peak before the other three, although there appears to be no clear ordering among the three. There is no discernable pattern in the peak 2 times among this set of regions in **D**.

#### Peak time correlations among the cortical areas tend to be stronger than those between cortical areas and LGN

We obtained the trial-to-trial correlations in peak times between pairs of areas from the entries of the matrix **R** described in the modeling section, and aggregated them across mice for both peaks using a weighted mean, where weighting is done using the standard error for each mouse. (As previously stated, this is a lower variance estimator than the simple arithmetic mean; see the discussion in (Kass et al., 2014, Chapter 8).) As shown in Figure 9, the cortico-thalamic correlations tended to be lower on average than the correlations among the cortical areas, although this was more striking for peak 1 than for peak 2.

**Figure 9:**
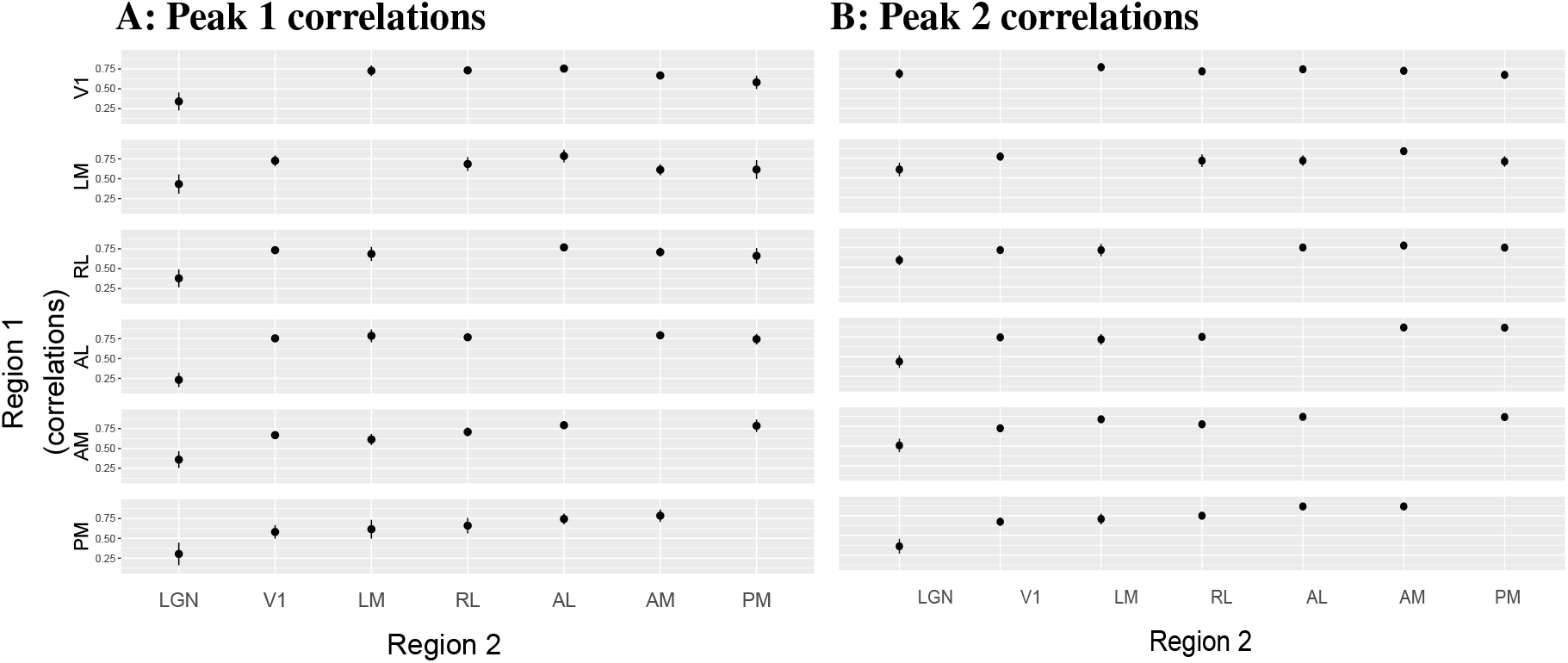
Trial-to-trial correlations in the peak times between pairs of areas. Each panel shows the weighted mean correlations of peak times between pairs of areas, across *N* = 13 mice, with their one standard error bars. Each row in each panel shows the correlations between a single visual area and all other areas. We observe in **A**, that the correlations in peak 1 times among the cortical areas tend to be stronger than those between cortical areas and LGN. We observe this to a lesser degree in **B**, along with the fact that peak 2 correlations tend to be stronger than their corresponding peak 1 correlations.

We also computed the mouse-to-mouse standard deviations of these correlations in each case. Figure 10 shows that the correlations between LGN and the cortical areas generally appear more variable across mice than correlations among cortical areas.

**Figure 10:**
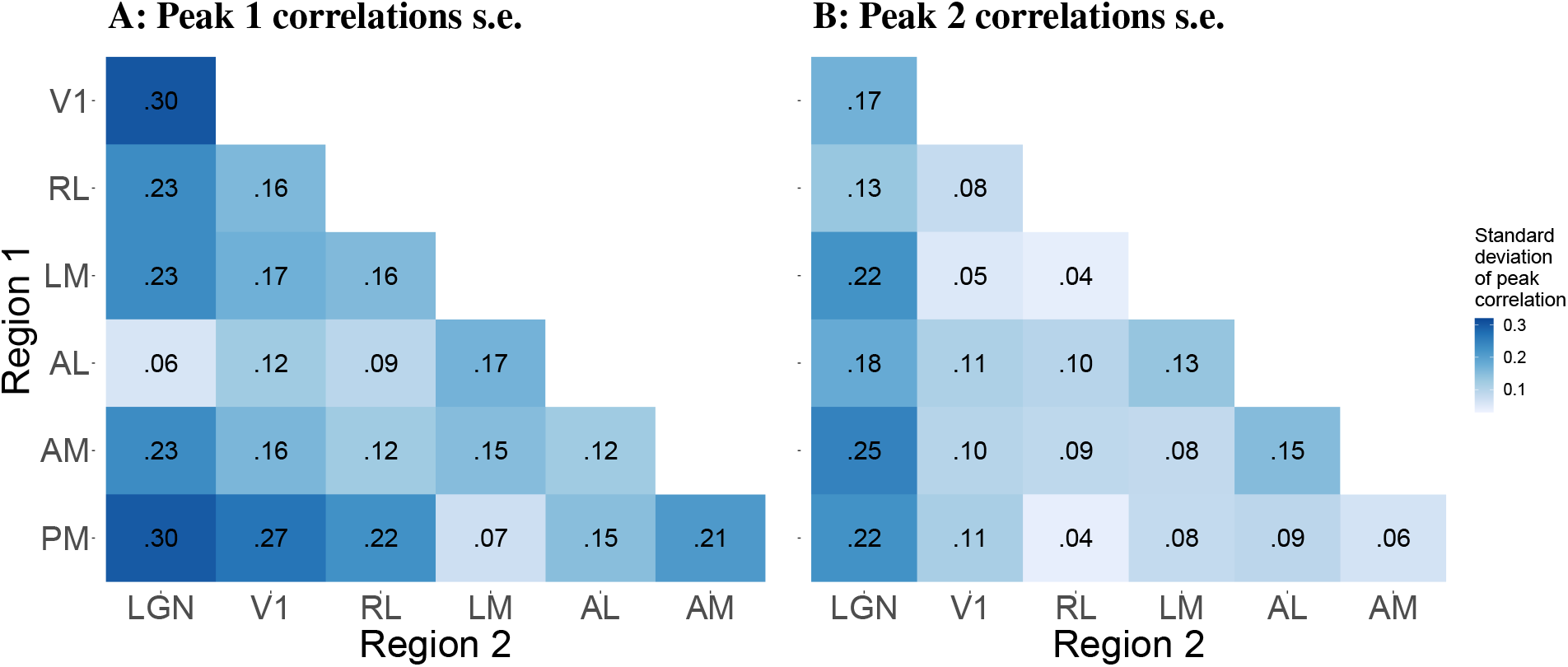
Standard deviations across mice of pairwise correlations between peak times. Each entry in the heat map represents one standard deviation of the pairwise correlations in peak times between the corresponding pair of regions. the color corresponds to the magnitude of the standard deviation. The figure reveals that the peak 1 correlations tend to be more variable across mice than the peak 2 correlations.

#### Correlations between thalamic and early cortical areas show the largest percentage changes after conditioning on V1

Given the neuroanatomy of the mouse thalamocortical pathway, we sought to quantify the involvement of V1 in the interactions between pairs of areas. We computed the partial correlations between the peak times for each pair of visual areas, conditioned on V1, and compared this to the original (marginal) correlation for the pair. Specifically, we computed the percentage decrease from marginal correlation to partial correlation for each pair of areas, where the partial correlation was conditioned on V1. As seen in Figure 11, for both peaks, the pair AM–PM has the least decrease in correlation given V1, which is expected given that they are farthest areas from V1 in the visual hierarchy, and there are no expected projections between this region pair through V1 Harris et al. (2019); Siegle et al. (2021). Among the cortical regions, according to the results for peak 1, the further downstream the region is from V1, the smaller the drop in its correlations with other regions after conditioning on V1. The biggest drops in correlations are seen in the correlations of LGN and the cortical areas, suggesting strong mediation of these interactions by V1. In particular, the correlation of LGN with RL for peak 1, goes from a positive to a negative correlation after conditioning on V1 (the decrease in correlation is greater than 100%). For peak 2 there are similar large decreases for pairs involving LGN, but the combination of feedback projection with the inconsistent timing results in Figure 8 (especially for LGN and V1) suggests the reversal of correlation between LGN and each of AM, AL, and PM after conditioning on V1 may be due to bidirectional connections between V1 and AM, AL, and PM.

**Figure 11:**
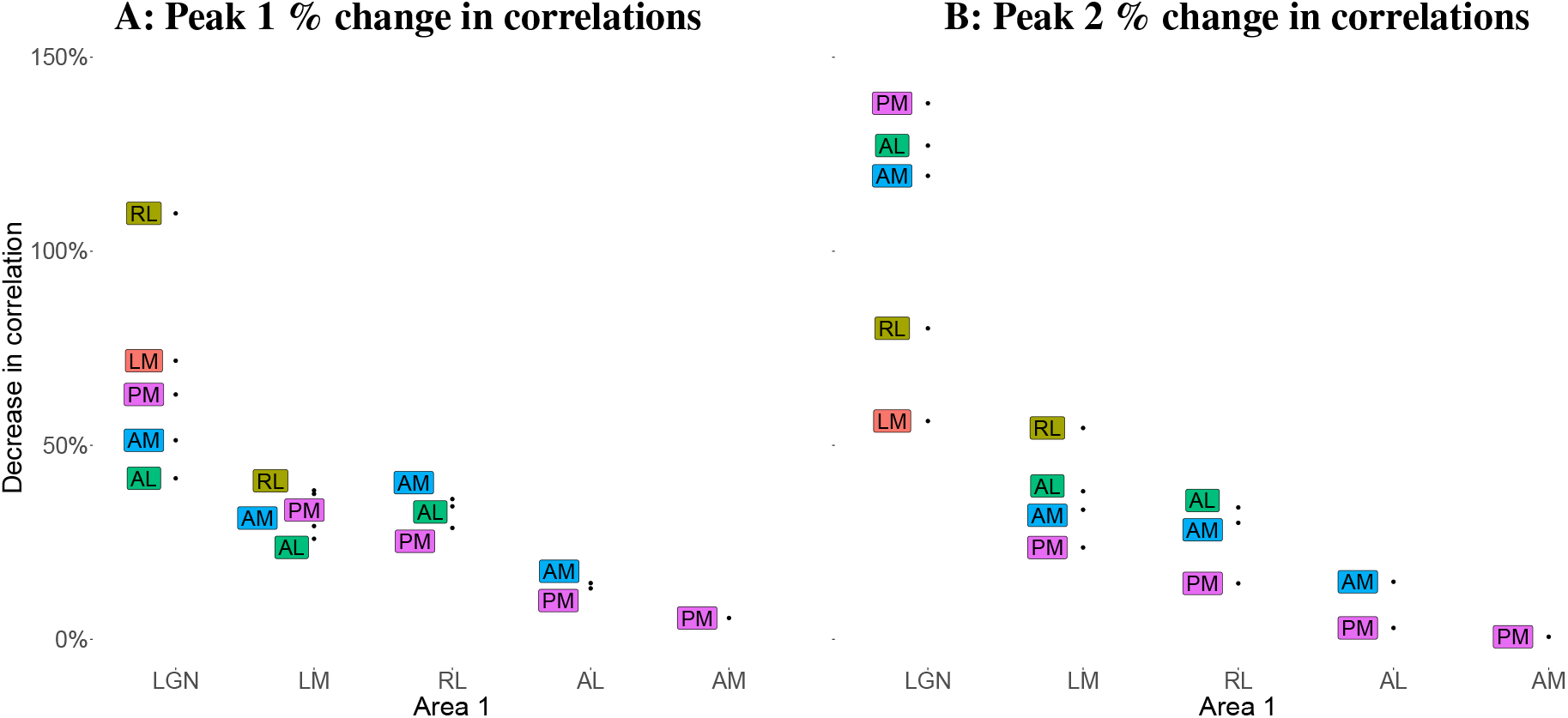
Percentage decrease in correlations between pairs of areas after conditioning on V1. Each labeled point shows the percentage decrease in the peak time correlations for the region pair consisting of the text label region and the corresponding region on the x-axis. For example, after conditioning on V1, the correlation between the peak 1 times in LGN and AL decreased by about 38%. The standard errors in all cases are *<* 10%. The previously positive correlation between LGN and RL in peak 1, and between LGN and AM, AL and PM in peak 2 became negative after conditioning V1 (the decrease in correlation is greater than 100%).

## Discussion

Studies of sequential timing of activity across brain areas have generally relied on data aggregated across trials. We aimed to develop, assess, and illustrate a relatively simple and computationally efficient method for identifying precise trial-by-trial sequential timing in population activity across areas. Motivated by results of Chen et al. (2022), we created a straightforward 3-step procedure and found it was nearly as accurate as the more complicated methodology in Chen et al. (2022) while running about 10 times faster, which enabled our comparative analysis of data from multiple subjects. This powerful method is accessible to the many neuroscientists who could apply it to recordings from large populations of spiking neurons.

Our examination of the variability in sequential timing relationships across 13 mice in the Allen Brain Observatory Visual Coding Neuropixels dataset Allen Institute MindScope Program (2019) produced results that are consistent with known anatomy and physiology, while also highlighting the distinction between pathways that are consistent across subjects versus those that are idiosyncratic. In the feedforward case of peak 1, for example, while LGN activity always precedes V1 acticity which always precedes activity in higher-order visual areas, most of the timing relationships among those higher-order visual areas are subject dependent. In the case of peak 2, which involves feedforward, feedback, and inputs from other areas, the relative timing of LGN and V1 is subject dependent.

We note that the subject-to-subject variability we observed may be attributed primarily to differences in animal physiology, differences in the experimental setup used in data collection, or to a combination of both these and other factors. Although inconsistencies across mice remain to be explained in greater detail, the results that were nearly the same across mice are compatible with existing notions of a functionally relevant hierarchical visual pathway for feedforward signal propagation. One interesting set of results that await further exploration involves LGN: the correlations in peak time between LGN and the cortical areas are weaker, on average, and more variable across mice, than those among the visual cortical regions themselves.

A potential explanation is that targeting Neuropixels probes to deep brain structures such as LGN, based solely on a map of the visual cortex, is prone to inaccurate placement, resulting in poor representation of relevant LGN neural populations and increased variability across mice. Similarly, partial correlations after conditioning on populations that are incompletely sampled by electrodes must be interpreted carefully. The contrast of the substantial decrease in correlation between LM and AM, after conditioning on V1, versus the small decrease in correlation between PM and AM may seem consistent with notions of visual hierarchy. On the other hand, the modest decrease in correlation between LGN and PM, after conditioning on V1 might be due to the many paths from V1 to PM (which could create variation in timing, as seen in the reduced correlation of V1 with PM compared to other areas) or it could be that key projection neurons may not have been sampled.

To the extent that the substantial mouse-to-mouse variability in functional connectivity observed across particular visual areas may have physiological sources, they could be genetic, developmental, or experiential factors (or combinations of these). Future studies could explore these distinctions, for example by comparing response timing in different mouse lines or mice with different types of visual exposure (e.g. dark-reared). Such investigations would be especially useful if combined with causal manipulations.

We have demonstrated the usefulness of this method in estimating peak times and their trial-to-trial correlations for spike train data. Although it is notoriously difficult to tease out informative trial-to-trial fluctuations in continuous data such as EEGs, Klein *et al*. Klein et al. (2021) decomposed local field potentials (LFPs) into current source densities (CSDs) on a trial-by-trial basis from which they demonstrated cross-population frequency coupling that was not apparent from the original LFPs. A variant of the methodology developed here might, in a similar vein, be useful for establishing timing relationships from CSDs. On the other hand, our analysis of bursts in temporally-evolving firing rate functions takes advantage of the substantial information about the timing of their maxima; by definition, that is where the most spikes occur. We also strengthened covariation relationships by confining attention to a single, homogeneous sub-population of neurons in each area. By instead examining multiple sub-populations, future work could investigate the diversity of functional interactions across areas.

## Acknowledgements

This work was supported by the National Institute of Mental Health, RO1 MH064537, and by the Allen Institute. We thank the Allen Institute founder, Paul G. Allen, for his vision, encouragement, and support.

## Notes

**Conflict of interest statement:** The authors declare no competing financial interests.

https://allensdk.readthedocs.io/en/latest/visual_coding_neuropixels.html

### Competing Interest Statement

The authors have declared no competing interest.

